# Effect of paclitaxel treatment on cellular mechanics and morphology of human oesophageal squamous cell carcinoma in 2D and 3D environments

**DOI:** 10.1101/2022.03.06.483167

**Authors:** Martin Kiwanuka, Ghodeejah Higgins, Silindile Ngcobo, Juliet Nagawa, Dirk M. Lang, Muhammad H. Zaman, Neil H. Davies, Thomas Franz

**Author notes:** Corresponding author, Department of Human Biology, Faculty of Health Sciences, University of Cape Town, Private Bag X3, Observatory 7935.

## Abstract

During chemotherapy, structural and mechanical changes in malignant cells have been observed in several cancers, including leukaemia and pancreatic and prostate cancer. Such cellular changes may act as physical biomarkers for chemoresistance and cancer recurrence. This study aimed to determine how exposure to paclitaxel affects the intracellular stiffness of human oesophageal cancer of South African origin *in vitro*. A human oesophageal squamous cell carcinoma cell line WHCO1 was cultured on glass substrates (2D) and in collagen gels (3D) and exposed to paclitaxel for up to 48 hours. Cellular morphology and stiffness were assessed with confocal microscopy, visually aided morpho-phenotyping image recognition, and mitochondrial particle tracking microrheology at 24 and 48 hours. In the 2D environment, the intracellular stiffness was higher for the paclitaxel-treated than for untreated cells at 24 and 48 hours. In the 3D environment, the paclitaxel-treated cells were stiffer than the untreated cells at 24 hours, but no statistically significant differences in stiffness were observed at 48 hours. In 2D, paclitaxel-treated cells were significantly larger at 24 and 48 hours and more circular at 24 but not at 48 hours than the untreated controls. In 3D, there were no significant morphological differences between treated and untreated cells. The distribution of cell shapes was not statistically significant different across the different treatment conditions in 2D and 3D environments. Future studies with patient-derived primary cancer cells and prolonged drug exposure will help identify physical cellular biomarkers to detect chemoresistance onset and assess therapy effectiveness in oesophageal cancer patients.

**Insight:** Mechanical changes in cancer cells by chemotherapeutic drugs exposure offer possible physical biomarkers for chemoresistance and cancer recurrence. This study on human oesophageal squamous cell carcinoma indicated that *in vitro* paclitaxel treatment induced stiffening and enlarging of malignant cells in two-dimensional environments at 24 and 48 hours. In physiologically more relevant three-dimensional collagen matrices the paclitaxel treatment led to cellular stiffening at 24 hours but softening thereafter, without significant changes in cellular size at any time. The outcomes need to be confirmed in future studies with prolonged drug exposure and patient-derived primary cancer cells.

## 1. Introduction

In 2018, oesophageal cancer (OC) ranked as the seventh most pervasive and the sixth deadliest cancer, with about 579,000 (3.2% of 18.1 million) cancer incidences and 508,800 (5.3% of 9.6 million) deaths reported globally [1]. OC is categorised into oesophageal squamous cell carcinoma (OSCC) and oesophageal adenocarcinoma (OAC), the two most prevalent histological subgroups. OSCC originates from the flat cells lining the upper two-thirds of the oesophagus, whereas OAC occurs in the lower third of the oesophagus and generally develops from Barrett mucosa [2]. OAC is most prevalent in high-income countries. In contrast, OSCC is the most prevalent subtype in South-East and Central Asia and Southern and Eastern Africa and is responsible for approximately 90% of all oesophageal cancer cases [2].

Oesophageal cancer is typically asymptomatic initially, and most patients present at a late stage when their chances of survival are dismal. As a result, OC has poor treatment outcomes, with a five-year survival rate of about 15% to 25% [3]. OC treatment options include endoscopic or surgical therapy and perioperative chemoradiotherapy [4]. Chemotherapy or chemoradiotherapy are indicated for patients with locally advanced OC or metastases in the lymph nodes. The chemotherapeutic drugs for OC treatment include carboplatin [5], paclitaxel [5, 6], cisplatin [6, 7], fluorouracil [7], and oxaliplatin [8]. The clinical effectiveness of chemoradiation therapy are associated with treatment failure, mainly due to radiation and chemoresistance [9, 10], the leading causes of OC recurrence [11, 12]. Therefore, there is a need to monitor chemoradiotherapy effectiveness and detect radiation resistance, chemoresistance and OC recurrence.

Exposure to chemotherapeutic drugs has been reported to increase cell stiffness in several cancers such as leukaemia [13], pancreatic cancer [14] and prostate cancer [15]. Cellular stiffening has been ascribed to cytoskeletal changes during exposure to chemotherapeutic drugs that aim to induce cell death, for example, by inhibiting cytoskeletal reorganisation required for cell division and proliferation [16–18].

Structural and mechanical changes in cancer cells associated with chemotherapy offer possible physical biomarkers to detect chemoresistance and cancer recurrence. However, there are no studies on the effect of chemotherapeutic treatment on cell mechanics of oesophageal cancer.

Hence, this study aimed to determine how exposure to the chemotherapeutic drug paclitaxel *in vitro* changes the intracellular stiffness and morphology of human oesophageal squamous cell carcinoma of South African origin in two-dimensional (2D) and three-dimensional (3D) environments. The study is based on the hypothesis that cancer cells within a tumour can respond differently to the same chemotherapeutic drugs by acquiring diverse strategies such as alterations in intracellular stiffness, morphologies, and organisation of the cytoskeleton, which decrease the efficacy of chemotherapeutic drugs leading to chemoresistance overtime.

## 2. Materials and methods

### 2.1 Cell culture

A human oesophageal squamous cell carcinoma (OSCC) cell line, WHCO1, derived from biopsies of South African patients [19], was used in this study. Cells were maintained in 25 cm^2^ tissue culture flasks (Coster, Corning Life Science, Acton, MA) in a humidified incubator at 37°C and 5% CO_2_. The cells were cultured in DMEM (Dulbecco’s Modified Eagle Medium, SIGMA, Life Science, USA) supplemented with 10% foetal bovine serum (FBS, Gibco, Life Technologies) and 1% (v/v) streptomycin-penicillin stock (Gibco, Life Technologies).

### 2.2 Chemotherapeutic drug treatment

For the 2D experiments, cells were grown on 35 mm glass-bottom dishes (Greiner Bio-One, Germany) for 24 hours and kept in a humidified incubator at 37°C and 5% CO_2_. The cells were then treated with 2 μM of paclitaxel (Teva Pharmaceuticals Pty Ltd) in fresh growth media for up to 48 hours.

For the 3D experiments, cells were grown as single cells in 3D collagen I gel at a final concentration of 2 mg/ml and treated with paclitaxel for up to 48 hours. Briefly, 1.4 x 10^5^ cells/10 μl were combined with equal amounts of Type I Rat Tail collagen (BD Bioscience, San Jose, CA) and neutralising solution (100 mM HEPES dissolved in 2xPBS, pH 7.4, SIGMA Life Science), and further diluted in 1xPBS until the desired 2 mg/ml collagen concentration was reached. A collagen volume of 50 μl was then gelled on 35 mm glass-bottom dishes kept in a humidified incubator at 37°C and 5% CO_2_ for an hour. After complete collagen gelation, the embedded cells were treated with 2 μM of paclitaxel in fresh growth media for up to 48 hours.

Vehicle control experiments with ethanol alone (0.014% [v/v], maximum volume used in the drug treatment experiments) were carried out for the 2D experiments to rule out the effect of ethanol on the cells. The ethanol-treated cells were found to have similar mean squared displacement (MSD) and power-law coefficient α curves compared with the untreated cells. Ethanol treatment was therefore not performed in the 3D experiments.

### 2.3 Mitochondrial particle tracking microrheology

After the 24 and 48 hours of paclitaxel treatment, MitoTracker Green solution (Life Technologies, Carlsbad, CA) was incubated with the cells for 30 minutes (2D) and 45 minutes (3D) at 37°C and 5% CO_2_ before the microrheology experiments. 100 nM and 400 nM of MitoTracker green solution in fresh growth media were used to fluorescently label the mitochondria for the cells in 2D and 3D matrices, respectively. Using endogenous and abundant mitochondria as tracer particles has been validated for short delay times against exogenously injected nanoparticles [20].

#### 2.3.1 Time-lapse image acquisition

Time-lapse images were acquired for the cells that survived paclitaxel treatment using a confocal microscope (Zeiss LSM 880 Airy-scan, Germany) with a monochrome charge-coupled device (CCD) camera and a 63 X 1.4 NA oil immersion objective. Cells were maintained in the incubation chamber of the confocal microscope at 37°C and 5% CO_2_ throughout the imaging sessions. ZEN software was used to capture the time-lapse images. During the imaging sessions, single cells were located and tracked for 140 and 150 seconds (approximately seven frames per second) for cells in 2D and 3D, respectively.

For the cells in 3D, only single cells near the centre of the gel were imaged to eliminate the boundary effects. No large focal drifts were observed as individual cells were imaged for only a maximum of 150 seconds. Time-lapse images were captured for at least ten individual cells in each condition. Cells that seemed to be dividing were eliminated from the imaging experiments and image analysis.

#### 2.3.2 Image processing and analysis

For cells containing at least 75 mitochondria, the particle trajectories describing mitochondrial fluctuations within the cytoplasm were extracted from the time-lapse images for each cell using TrackMate, a Fiji Image J plugin [21, 22]. The particle trajectories were converted to time-dependent mean-squared displacement (MSD) using MATLAB (Math Works, Inc., 2020) and MSD-Analyzer class [22, 23] according to:

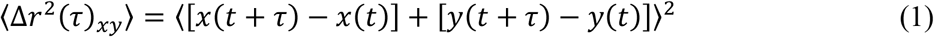

where 〈Δr^2^ (τ)_*xy*_〉 is the ensemble-averaged MSD, τ is the time interval or delay time between the first and last image frame analysed, and x(t) and y(t) are the spatial coordinates of a particle position at the time t.

For viscoelastic materials, the MSD scales nonlinearly with the delay time τ according to a power-law relationship, i.e. 〈Δr^2^ (τ)〉 ~ τ^α^ [24]. The power-law coefficient α = *∂*ln 〈Δr^2^ (τ)/〉ln (τ) represents the slope of the logarithmic MSD-τ curve.

Lower and flat MSD represents constrained particle fluctuations and stiffer, solid material, whereas a high and increasingly sloped MSD indicates greater particle motility and a more fluid-like material.

The mitochondrial MSD primarily represents the viscoelastic intracellular properties for short delay times, whereas active intracellular motor-driven processes dominate the MSD for long delay times [20, 25]. The power-law coefficient α helps classify the motion of the particles: α ≈ 1 corresponds to diffusive motion such as thermal fluctuations in Newtonian fluids, and α ≈ 0 represents constrained, sub-diffusive motion such as thermal fluctuations in an elastic material [26, 27]. For a non-equilibrium system such as the intracellular environment, molecular motor activity results in additional internal fluctuations beyond thermal fluctuations leading to increased MSD and α at relatively longer times scales [20, 28]. Hence, short delay times data (0 to 1 seconds) were used to extract the viscoelastic properties of cells.

### 2.4 Morphological analysis

Morphological analysis was performed using the visually aided morpho-phenotyping image recognition (VAMPIRE) algorithm [29, 30]. Briefly, brightfield images of the treated and untreated single cells (dividing cells and cells at the boundary of the gel in 3D were excluded from this study) in 2D and 3D were captured using a confocal microscope and then segmented using the Trainable Weka Segmentation plugin in Fiji Image J [31] to identify cell boundaries. The segmented images were imported to VAMPIRE and used to construct a shape analysis model (identify representative shape modes among all shape modes present in the cell population), which was applied to our dataset to analyse the shapes of treated and untreated cells. In addition to determining the fractional abundance of the treated and untreated cells in each shape mode, it was also used to determine the cells’ area, perimeter, circularity, and aspect ratio.

### 2.5 Data and statistical analysis

Statistical analysis was carried out using a two-way ANOVA unless otherwise stated. Residual analysis was used to evaluate the assumptions of a two-way ANOVA. Shapiro-Wilk and Levene’s tests were used to determine the normality of data and homogeneity of variance, respectively. Upon detection of statistical significance, post-hoc analysis was carried out using Tukey’s test. Statistical significance was assumed at *p* < .05. Data are presented as mean ± standard deviation unless indicated otherwise. Data are available in the online supplement. All statistical analyses were carried out using SPSS for Windows 26.0. (IBM Corp, Armonk, NY). All experiments were run in duplicate on different days.

## 3. Results

### 3.1 Intracellular stiffness in 2D environments

Mitochondrial particle tracking microrheology (MPTM) experiments were performed to determine the effect of paclitaxel treatment on the intracellular stiffness of cells in 2D environments. Figure 1 shows representative images of paclitaxel-treated, ethanol-treated, and untreated single cells at 24 and 48 hours. The green-stained mitochondria were used as the tracer particles for the MPTM experiments. The mitochondrial fluctuations within the cytoplasm of the cells were tracked for 26, 22 and 31 cells in the paclitaxel-treated, ethanol-treated, and untreated group, respectively.

**Figure 1:**
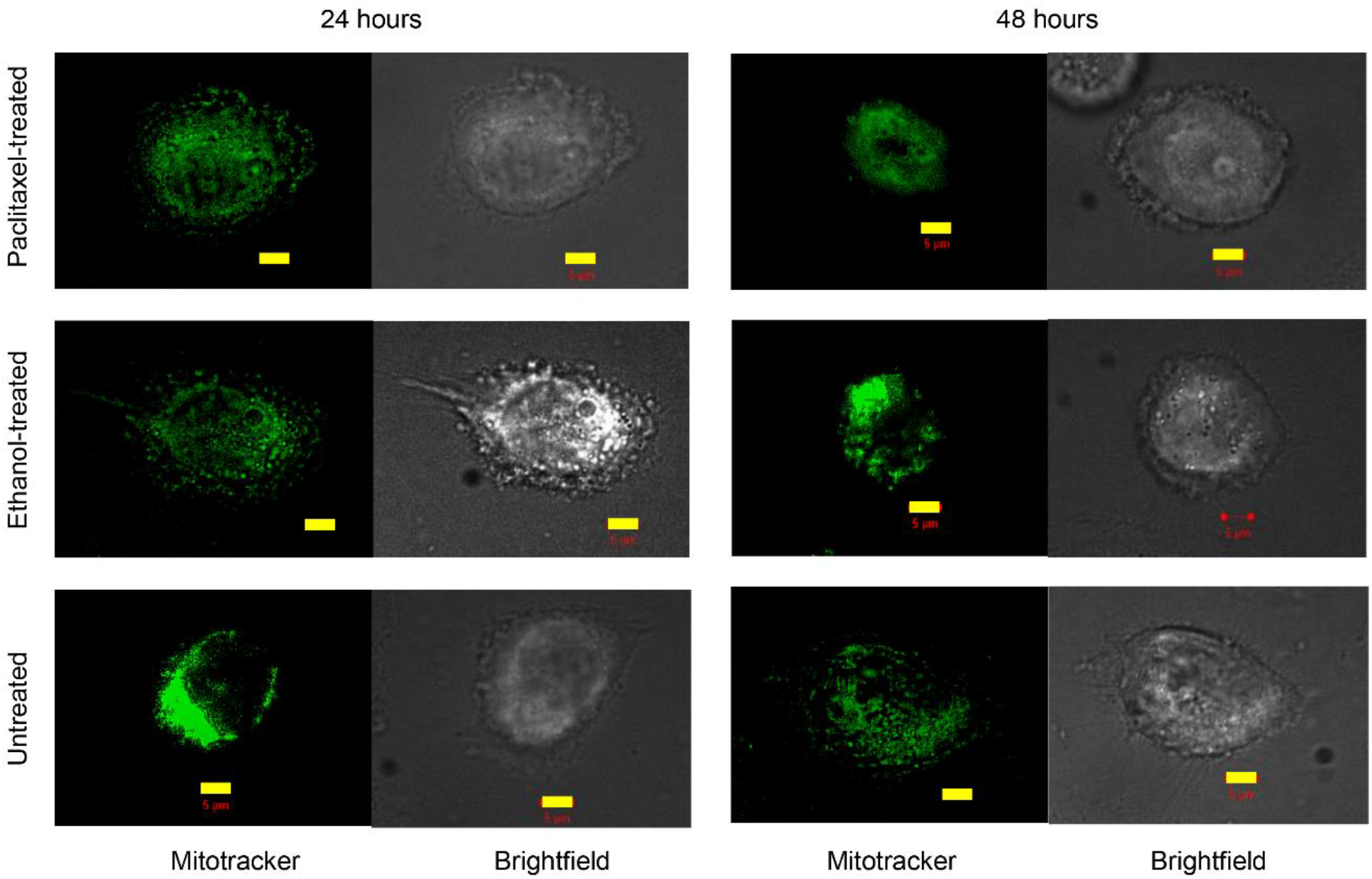
Representative fluorescence and brightfield images of paclitaxel-treated, ethanol-treated and untreated cells in 2D environments after 24 and 48 hours. Scale bars (yellow) represent 5 μm.

The paclitaxel-treated cells had significantly higher MSD at 24 hours than at 48 hours at all delay times (*p* < 0.05) except at τ = 0.14 seconds (Figure 2 a). Also, the power-law coefficient α of paclitaxel-treated cells was higher at 24 hours than at 48 hours for all delay times, with statistically significant differences only observed between τ = 0.28 and 2.1 seconds (*p* < 0.05) (Figure 2 b). There were no statistically significant differences in either MSD or α between 24 and 48 hours for ethanol-treated cells (Figure 2 c and d) and untreated cells (Figure 2 e and f).

**Figure 2:**
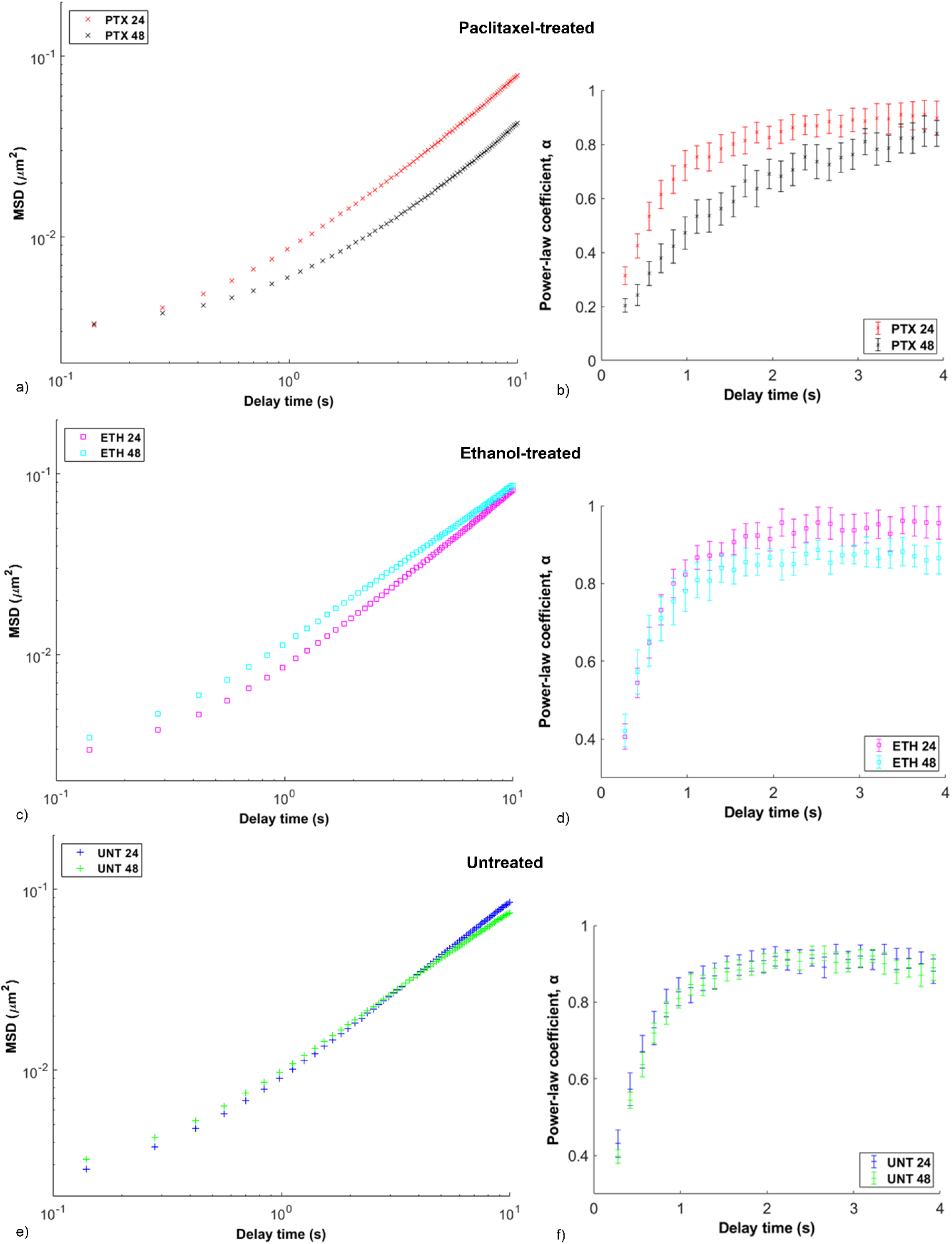
Time-point comparison of microrheological characteristics of cells in 2D environments. MSD versus delay time τ for paclitaxel-treated (PTX) (a), ethanol-treated (ETH) (c) and untreated cells (UNT) (e) at 24 and 48 hours. Power-law coefficient α versus delay time τ for paclitaxel-treated (b), ethanol-treated (d) and untreated cells (f) at 24 and 48 hours. Error bars are SEM for one independent repeat experiment (n = 2), N = 12 (PTX 24), 14 (PTX 48), 11 (ETH 24, ETH 48), 17 (UNT 24) and 14 (UNT 48). N refers to the number of cells in each condition.

At 24 hours of treatment, statistically significant differences in MSD were only observed between paclitaxel-treated and untreated cells at τ = 0.7 seconds (*p* = 0.031) (Figure 3 a). The power-law coefficient α of the paclitaxel-treated cells was lower than that of ethanol-treated and untreated cells at all delay times, with statistical significance only observed between paclitaxel-treated and untreated cells at τ = 0.28 seconds (*p* = 0.026) (Figure 3 b).

**Figure 3:**
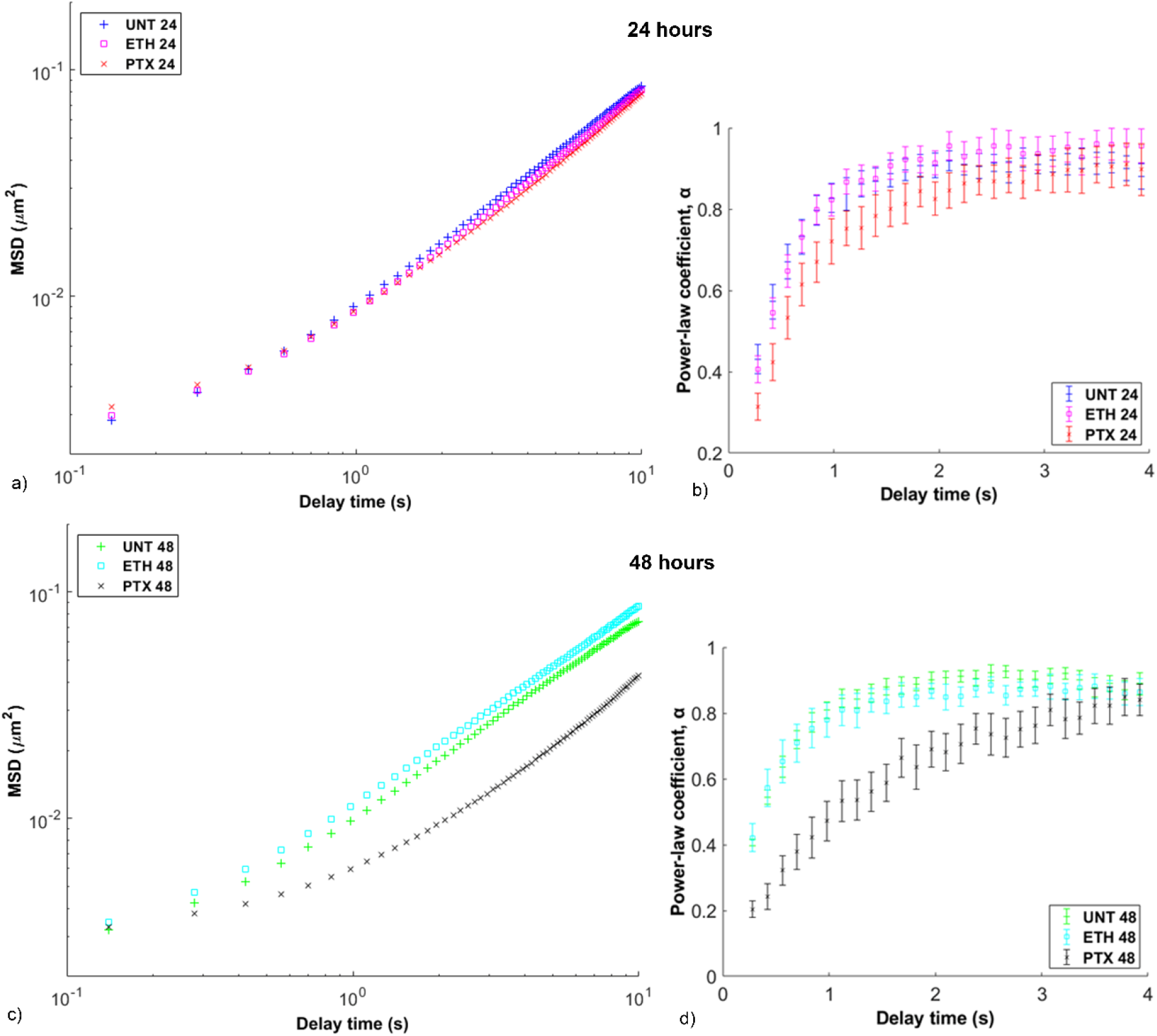
Treatment comparison of microrheological characteristics of cells in 2D environments. MSD versus delay time τ for paclitaxel-treated (PTX), ethanol-treated (ETH), and untreated cells (UNT) at 24 (a) and 48 hours (c). Power-law coefficient α versus delay time τ for paclitaxel-treated, ethanol-treated and untreated cells at 24 (b) and 48 hours (d). Error bars are SEM for one independent repeat experiment (n = 2), N = 17 (UNT 24), 11 (ETH 24), 12 (PTX 24), 14 (UNT 48), 11 (ETH 48) and 14 (PTX 48).

At 48 hours of treatment, the paclitaxel-treated cells had lower MSD than the ethanol-treated (*p* < 0.05) and untreated cells (*p* < 0.005) at all delay times except τ = 0.14 seconds. No statistically significant differences in MSD were observed between the ethanol-treated and untreated cells (Figure 3 c). The power-law coefficient α of the paclitaxel-treated cells was significantly lower than that of the ethanol-treated (*p* < 0.05) and untreated cells (*p* < 0.005), except for delay times between τ = 3.5 and 4 seconds. No statistically significant differences in power-law coefficient α were observed between the ethanol-treated and untreated cells (Figure 3 d).

### 3.2 Intracellular stiffness in 3D environments

Mitochondrial particle tracking microrheology experiments were performed to investigate whether paclitaxel treatment affected the intracellular stiffness of cells embedded as single cells in 3D matrices. Figure 4 shows representative images of the paclitaxel-treated cells and the untreated single cells at 24 and 48 hours of treatment with green-stained mitochondria as tracer particles for the MPTM experiments. The displacements of the mitochondria within the cytoplasm were tracked for a total of 31 paclitaxel-treated cells and 32 untreated cells.

**Figure 4:**
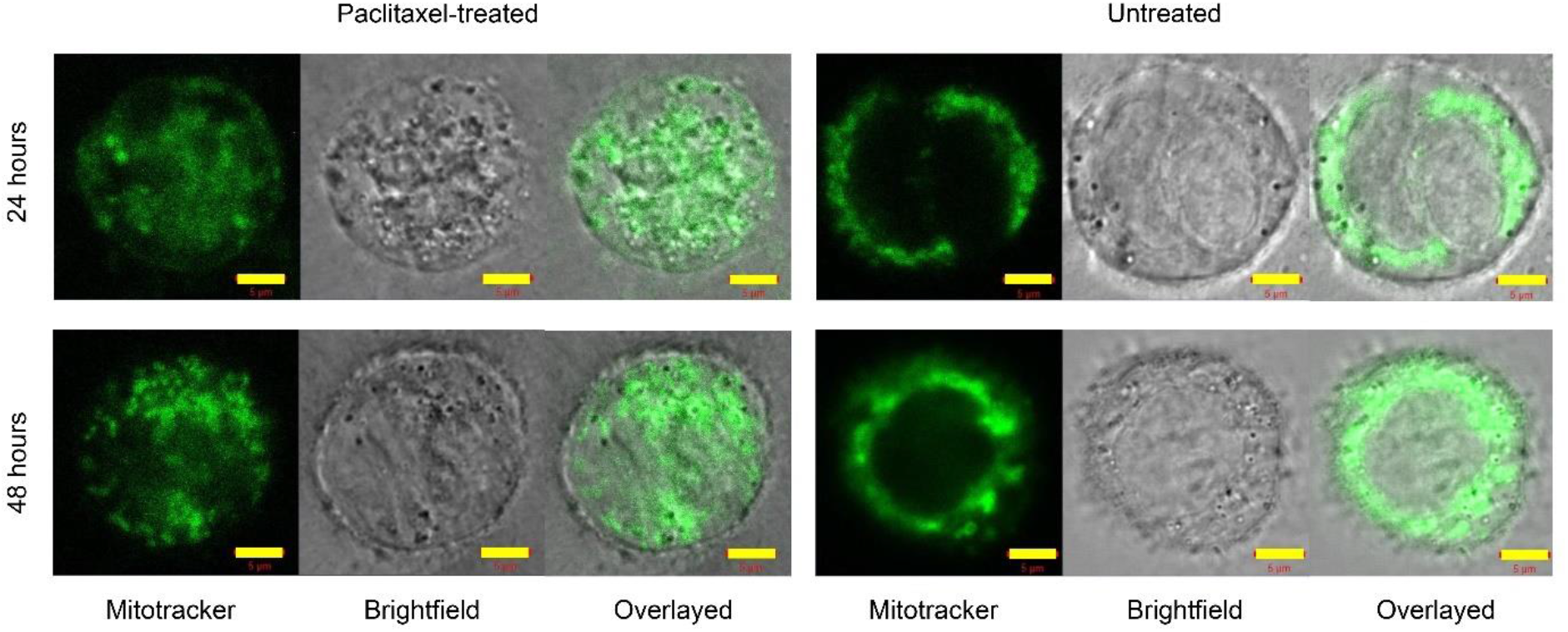
Representative fluorescence, brightfield, and overlay images of paclitaxel-treated, ethanol-treated and untreated single cells in 3D environments after 24 and 48 hours. Scale bars (yellow) represent 5 μm.

The paclitaxel-treated cells exhibited significantly lower MSD and α at 24 hours than at 48 hours for all delay times (*p* < 0.05) except τ = 0.15 seconds for MSD (*p* = 0.056) (Figure 5 a) and τ = 1.2, 1.5, and 2.1 seconds for α (Figure 5 b). The untreated cells exhibited higher MSD at 24 hours than at 48 hours at all delay times, with statistical significance observed between τ = 0.9 and 3.6 seconds (*p* < 0.05) (Figure 5c). Statistically significant differences in power-law coefficient α of the untreated cells between 24 and 48 hours were only observed between τ = 0.15 and 0.9 seconds (*p* < 0.05) (Figure 5d).

**Figure 5:**
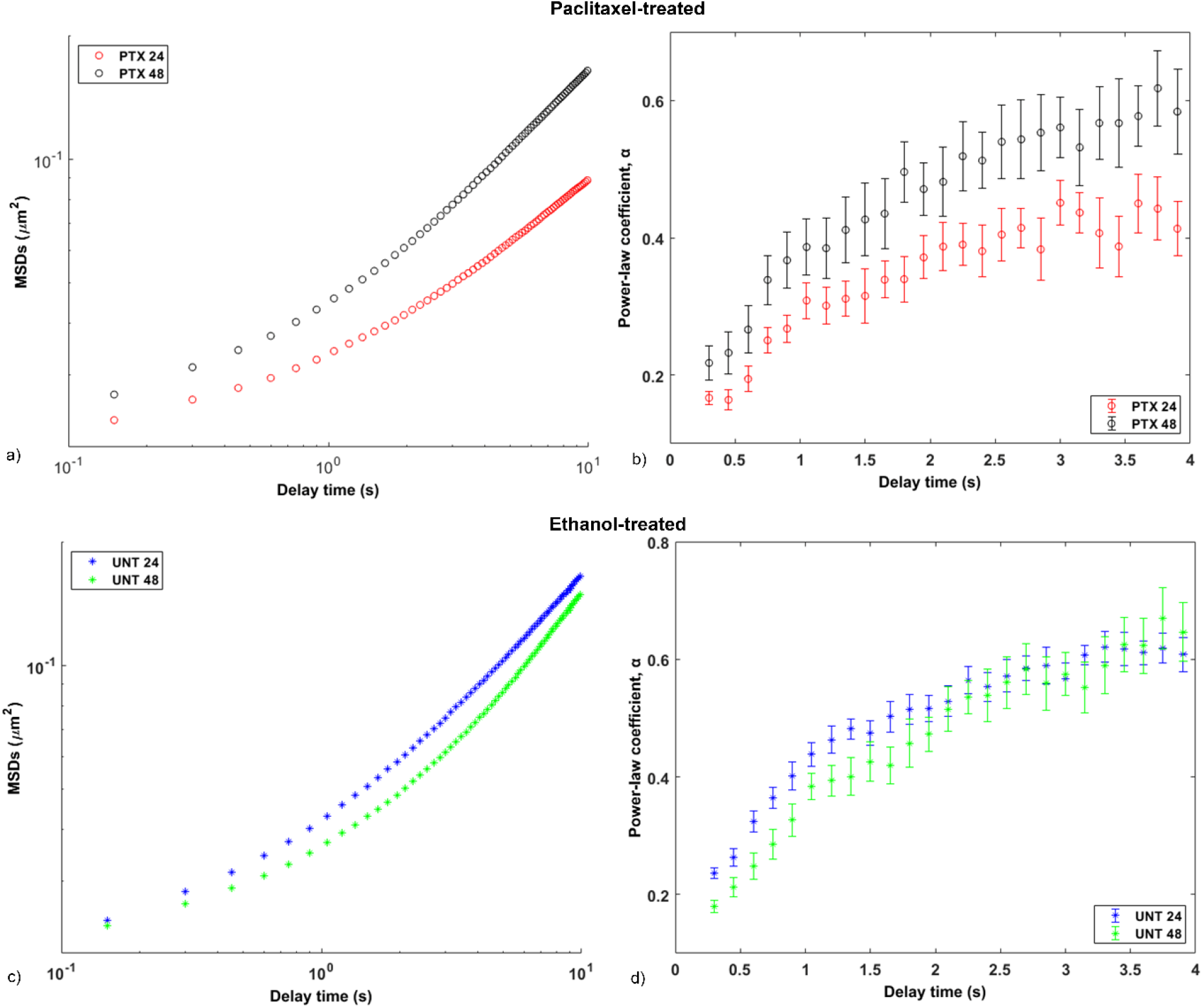
Time-point comparison of microrheological characteristics of cells in 3D environments. MSD versus delay time τ for (a) paclitaxel-treated cells at 24 (PTX 24) and 48 (PTX 48) hours, and (c) untreated cells at 24 (UNT 24) and 48 (UNT 48) hours. Power-law coefficient α versus delay time τ for (b) paclitaxel-treated cells at 24 and 48 hours, and (d) untreated cells at 24 and 48 hours. Error bars indicate SEM for one independent repeat experiment (n = 2); N = 21 (PTX 24), 10 (PTX 48), 19 (UNT 24) and 13 (UNT 48).

At 24 hours of treatment, the paclitaxel-treated cells had significantly lower MSD than the untreated cells at all delay times (*p* < 0.05) except between τ = 0.15 and 0.3 seconds (Figure 6 a), and α was discernibly lower for the paclitaxel-treated cells than the untreated cells at all delay times (*p* < 0.05) (Figure 6 b).

**Figure 6:**
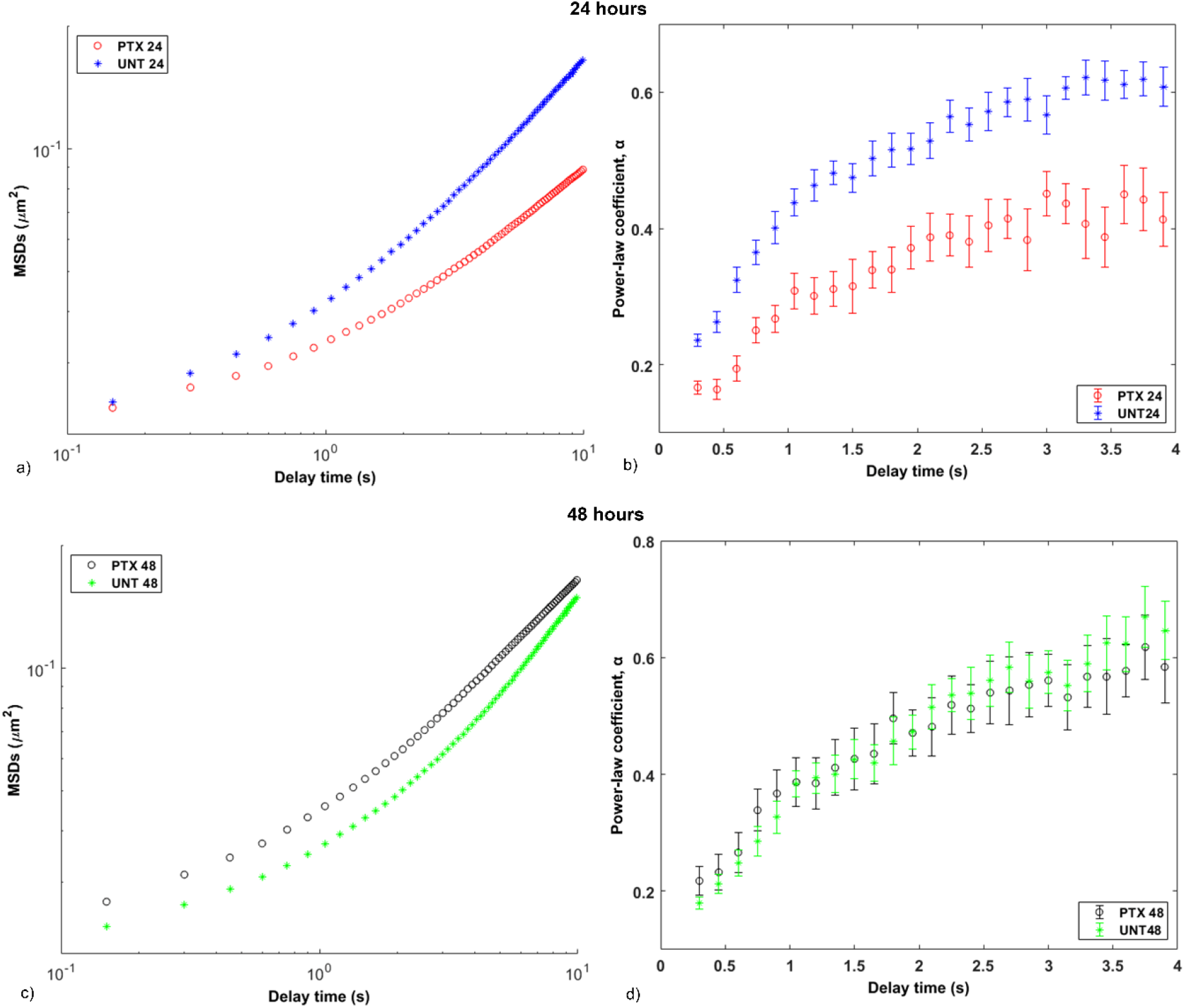
Treatment comparison of microrheological characteristics of cells in 3D environments. MSD versus delay time τ for paclitaxel-treated (PTX) and untreated cells (UNT) for 24 (a) and 48 hours (c). Power-law coefficient α versus delay time τ for paclitaxel-treated and untreated cells for 24 (b) and 48 hours (d). Error bars are SEM for one independent repeat experiment (n = 2), N = 19 (UNT 24), 21 (PTX 24), 13 (UNT 48) and 10 (PTX 48).

At 48 hours of treatment, the paclitaxel-treated cells exhibited higher MSD than the untreated cells at all delay times, with statistical significance between τ = 0.3 and 4.2 seconds (*p* < 0.05) (Figure 6 c). However, no statistically significant differences in power-law coefficient α were observed between the paclitaxel-treated and untreated cells (Figure 6 d).

### 3.3 Cell morphology

Cell area, perimeter, circularity, aspect ratio, and cell shape were analysed for 251 treated and untreated single cells using VAMPIRE. Of these, 194 cells were in 2D (49 paclitaxel-treated, and 66 ethanol-treated and 79 untreated cells), and 57 cells were in 3D matrices (26 paclitaxel-treated and 31 untreated cells). A three-way ANOVA was performed to assess the differences between groups for the various morphological parameters. Statistical significance was assumed at *p* ≤ 0.025.

In 2D environments, the paclitaxel-treated cells had a significantly larger cell area than the untreated (*p* < 0.005) and ethanol-treated cells (*p* < 0.0005) at both 24 and 48 hours (Figure 7 a). The paclitaxel-treated cells had a significantly larger perimeter than the ethanol-treated cells at 24 hours (*p* < 0.005) and the untreated cells at 48 hours (*p* < 0.005). The paclitaxel-treated cells also exhibited a significantly larger perimeter at 48 hours than at 24 hours (*p* < 0.005) (Figure 7 b). The paclitaxel-treated (*p* < 0.005) and ethanol-treated cells (*p* < 0.0005) were significantly more circular at 24 hours than 48 hours, whereas the opposite was observed for the untreated cells (*p* < 0.005). At 24 hours of treatment, the paclitaxel-treated cells were significantly more circular than the untreated and ethanol-treated cells (*p* < 0.0005). At 48 hours, no significant differences in circularity were detected between the groups (Figure 7 c). No significant differences in aspect ratio were detected between groups (Figure 7 d).

**Figure 7:**
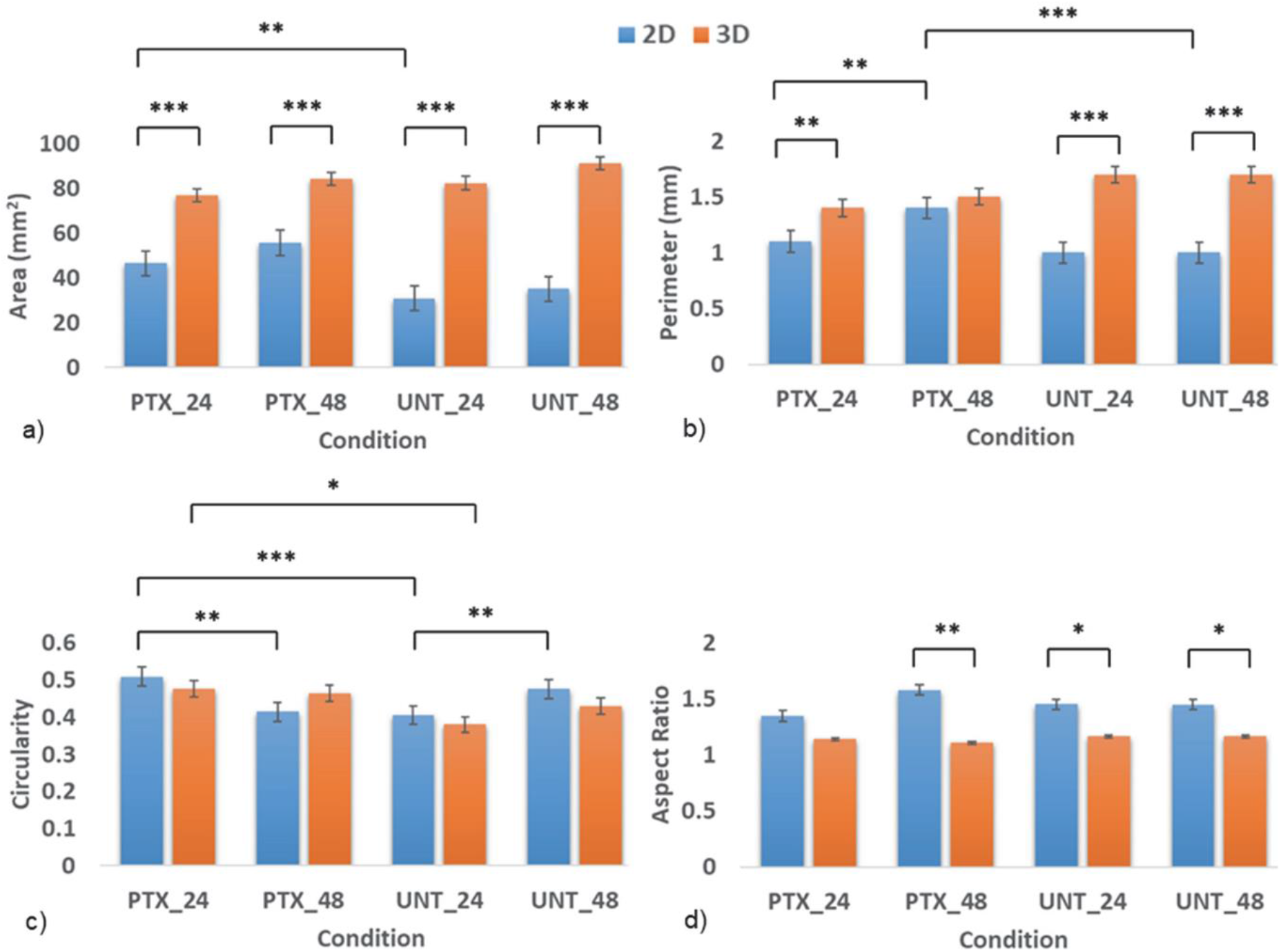
Morphometric data of paclitaxel-treated (PTX) and untreated cells (UNT) in 2D and 3D environments at 24 and 48 hours: Area (a), perimeter (b), circularity (c) and aspect ratio (d). Error bars are SEM for one independent repeat experiment (n = 2), N = 27 (2D) + 15 (3D) (PTX_24), 22 + 11 (PTX_48), 42 + 16 (UNT_24) and 37 + 15 (UNT_48). *, **, *** indicate p < 0.025, 0.005, and 0.0005, respectively.

In 3D matrices, no significant differences between groups were observed for cell area, perimeter and aspect ratio (Figure 7 a, b and d). Circularity was significantly higher in paclitaxel-treated than untreated cells at 24 hours (*p* < 0.025) (Figure 7 c).

Comparing between 2D and 3D environments, the paclitaxel-treated cells exhibited a significantly larger cell area in 3D than in 2D at both 24 and 48 hours (*p* < 0.0005), and a significantly larger perimeter at 24 hours (*p* < 0.005) but not at 48 hours (Figure 7 a and b). The untreated cells exhibited significantly larger cell area and perimeter in 3D than 2D at 24 and 48 hours (*p* < 0.0005). Cell circularity did not differ significantly between 2D and 3D environments for all treatment conditions and times (Figure 7 c). Significant differences in aspect ratio were only observed between the untreated cells in 2D and 3D at both 24 and 48 hours (*p* < 0.025) and the paclitaxel-treated cells in 2D and 3D at 48 hours (*p* < 0.005) (Figure 7 d).

Ten shape modes were identified from the entire data set of 251 cells using VAMPIRE. These modes represent the inherent shapes of the paclitaxel-treated and untreated cells in 2D and 3D matrices and ethanol-treated cells in 2D (Figure 8). Modes 1 and 2 represent an elongated and circular cell shape, respectively. The modes 3 through 10 represent irregular shapes. The shape modes were further categorised into five groups based on their similarity, as shown by the branches on the shape mode dendrogram (Figure 8 b). Shape modes 1 and 2 were categorised in groups 1 and 2, respectively. Shape modes 3, 4 and 5 are categorised in group 3, shape modes 6, 7 and 8 in group 4, and shape modes 9 and 10 in group 5. The Kruskal-Wallis test was used to test for differences between the distribution of the shape modes in the treatment groups for the 2D and 3D environments. The distribution of the shape mode groups among the paclitaxel-treated, ethanol-treated and untreated cells in 2D and 3D matrices were then used to assess the heterogeneity of cell shapes for the different conditions. The cells in a particular treatment condition were considered heterogeneous if their shape modes were well distributed in all the five shape mode groups.

**Figure 8:**
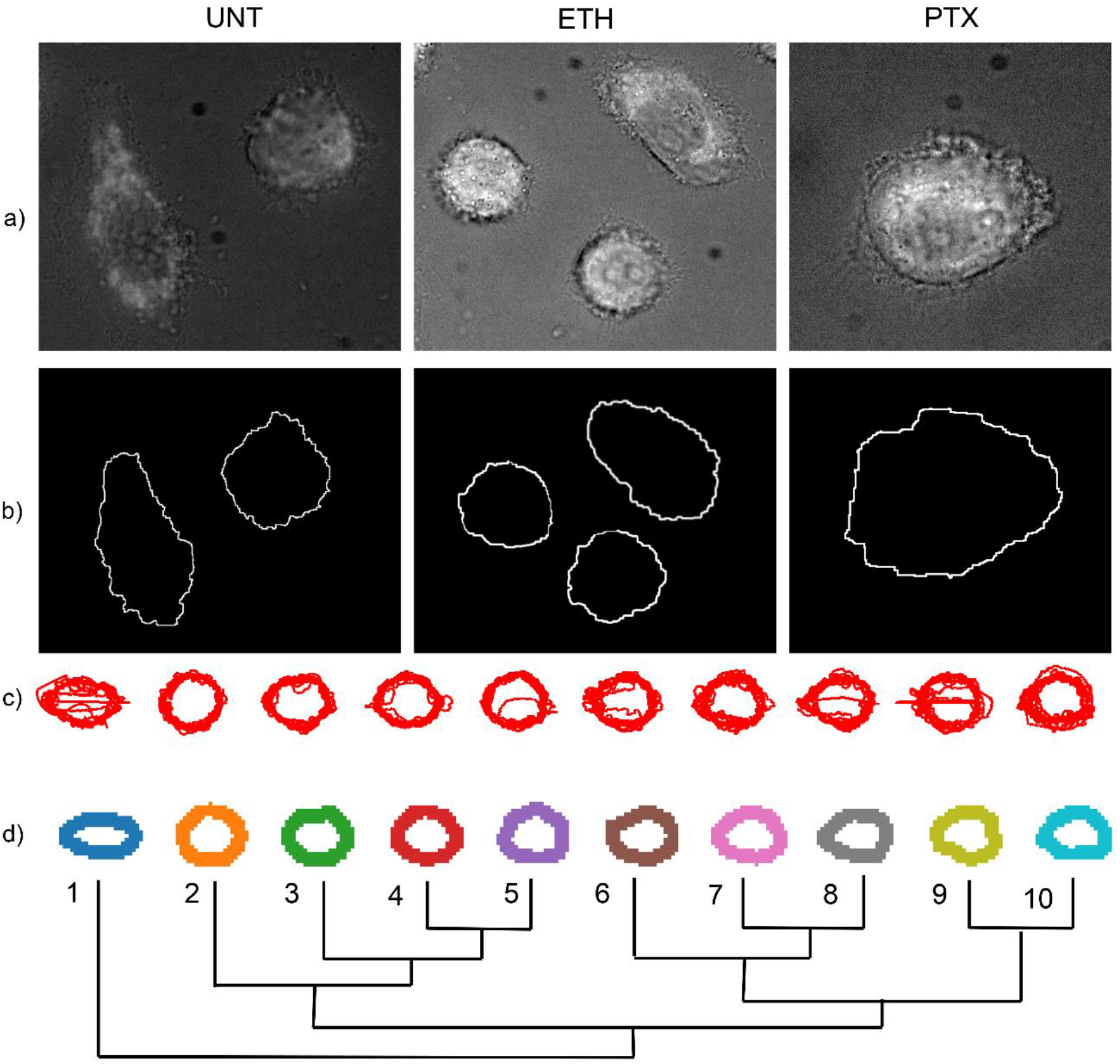
Shape modes obtained for WHCO1 cells in 2D and 3D environments using VAMPIRE. (a) Representative brightfield images untreated, ethanol-treated and paclitaxel-treated WHCO1 cells. (b) Representative images of cells segmented to identify boundaries that VAMPIRE algorithm uses to assign shape modes to individual cells in the population. (c) Registered objects show actual boundaries of cells selected randomly from individual shape modes. (d) Shape mode dendrogram demonstrates the different shape modes (1 through 10) detected and their similarity (shown by the branches).

For the 2D environment (Figure 9), the cellular shape modes in the different treatment conditions were well distributed in all groups of the shape modes. Results from the Kruskal-Wallis test showed that the distribution of the shape modes was significantly different across the different treatment conditions (*p* = 0.036, *η*^2^ = 0.0369; the effect size *η*^2^ is small). In the post-hoc analysis, pairwise comparisons between the distribution of shape modes for the different treatment groups were only significant for PTX 24 vs PTX 48 (*p* = 0.020) and ETH 24 vs UNT 24 (*p* = 0.044). However, considering Bonferroni correction, no significant differences (*p* > 0.097) were observed in the post-hoc analysis.

**Figure 9:**
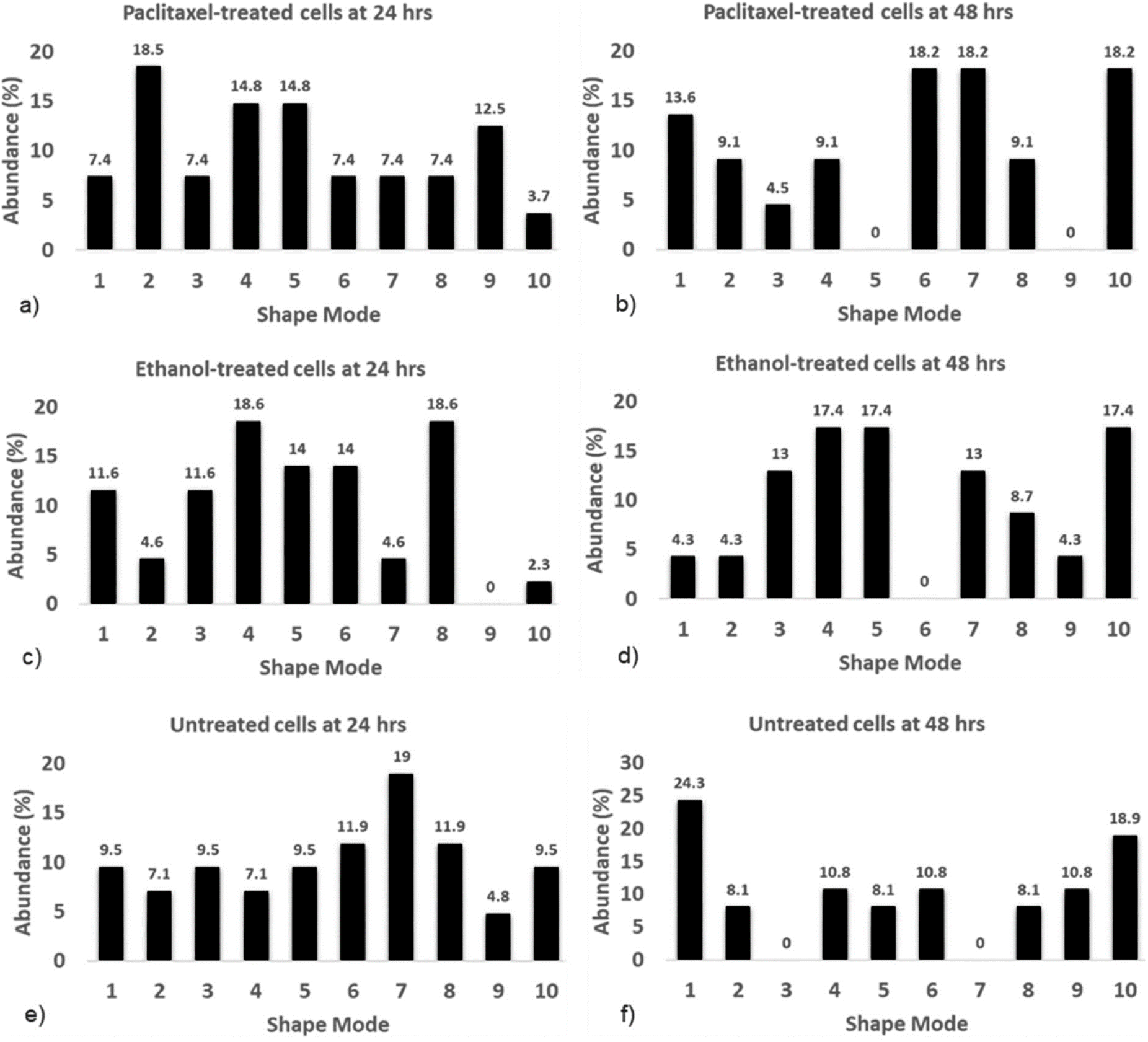
Distribution of shape modes in 2D environments. Paclitaxel-treated cells at 24 (a, N = 27) and 48 hours (b, N = 22), ethanol-treated cells at 24 (c, N = 43) and 48 hours (d, N = 23), untreated cells at 24 (e, N = 42) and 48 hours (f, N = 37). See Table S13 in the online supplement for absolute abundance, i.e. the number of cells assigned to each shape mode.

For the 3D environment (Figure 10), the cellular shape modes in the different treatment conditions were well distributed in all the shape mode groups except group 1 for paclitaxel-treated and untreated cells at 48 hours and the untreated cells at 24 hours. Results from the Kruskal-Wallis test showed that the distribution of the shape modes was not significantly different across the different treatment conditions (*p* = 0.086, *η*^2^ = 0.0677; the effect size *η*^2^ is moderate). In the post-hoc analysis, pairwise comparisons between the distribution of shape modes for the different treatment groups were only significant for PTX 48 versus UNT 48 (*p* = 0.033). However, considering Bonferroni correction, no significant differences (*p* > 0.163) were observed in the post-hoc analysis.

**Figure 10:**
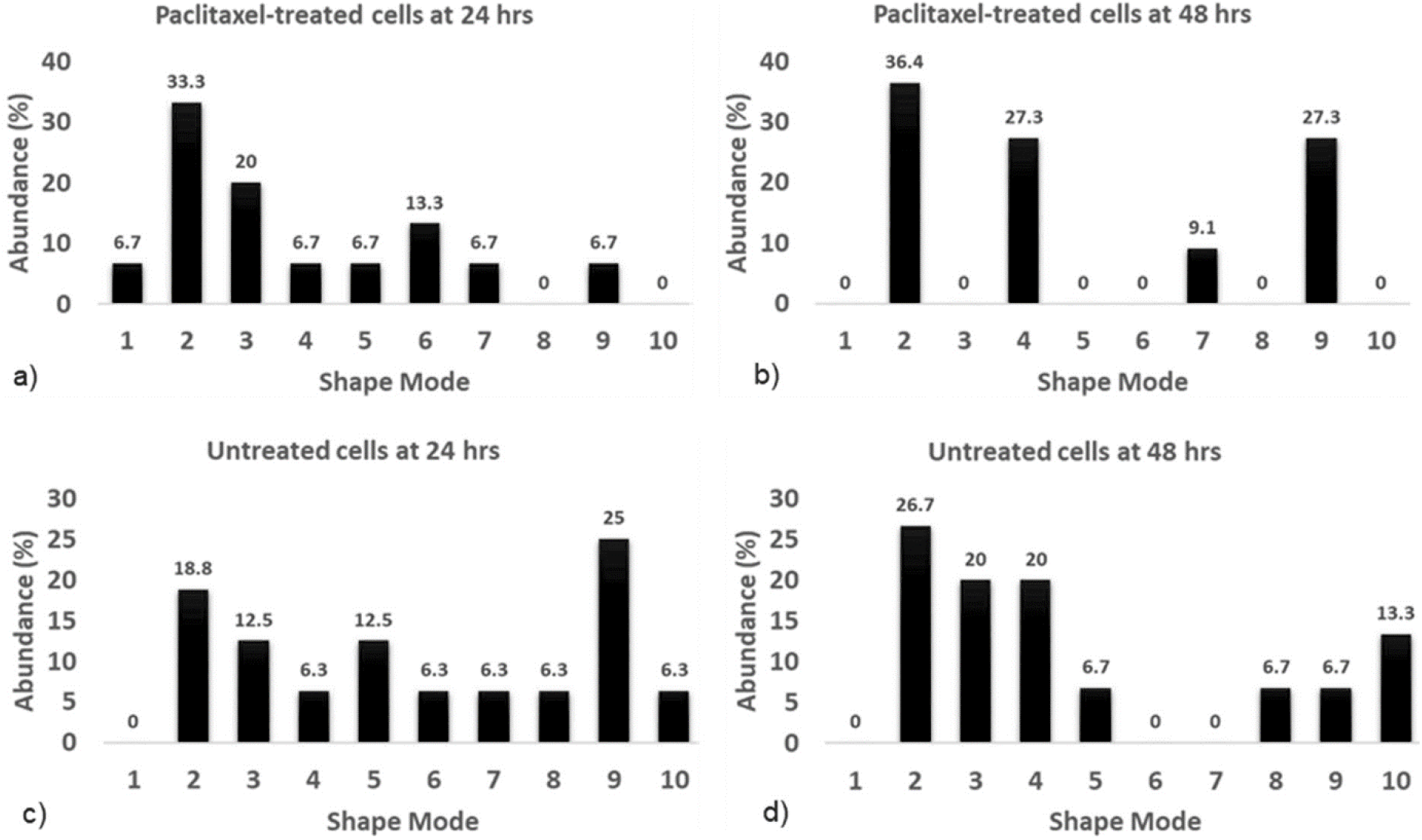
Distribution of shape modes in 3D environments. Paclitaxel-treated cells at 24 (a, N = 15) and 48 hours (b, N = 11), untreated cells at 24 (c, N = 16) and 48 hours (d, N = 15). Please see Table S13 in the online supplement for absolute abundance, i.e. the number of cells assigned to each shape mode.

On comparing the distribution of shape modes in 2D and 3D using the Kruskal-Wallis’ test, no significant differences were detected across the different treatment conditions at 24 hours (*p* = 0.446, *η*^2^ = 0.0035; the effect size *η*^2^ is small) and 48 hours (*p* = 0.142, *η*^2^ = 0.0301; the effect size *η*^2^ is small). In the post-hoc analysis, significant differences were only detected between the untreated cells at 48 hours (*p* = 0.043). However, considering Bonferroni correction, no significant differences (*p* > 0.261) were observed in the post-hoc analysis.

## 4. Discussion

### 4.1 Intracellular stiffness

This study characterised the effect of paclitaxel treatment on the intracellular stiffness and morphology of human OSCC cells of South African origin in 2D and 3D environments. The results discussed for intracellular stiffness are extracted from the power-law coefficient versus delay time plots for both cells in 2D and 3D matrices and are read from the delay times between 0 and 1 second. The paclitaxel-treated cells in 2D matrices exhibited higher intracellular stiffness (i.e. lower power-law coefficient α) than the untreated controls at both 24 and 48 hours of treatment. In addition, there was a significant increase in the intracellular stiffness of the paclitaxel-treated cells with an increase in drug exposure time from 24 to 48 hours. The absence of a significant stiffness increases with time for the untreated cells confirmed that the prolonged drug exposure led to the stiffening of the paclitaxel-treated cells.

When embedded in 3D collagen matrices, the paclitaxel-treated cells were significantly stiffer than the untreated cells at 24 hours but not 48 hours. The intracellular stiffness significantly decreased for paclitaxel-treated cells, whereas it significantly increased for the untreated cells from 24 to 48 hours. Thus, it is unclear whether and how intracellular stiffness changes in the paclitaxel-treated cells between 24 and 48 hours are related to drug exposure, as intracellular stiffening was also observed for the control at 48 hours.

Results on intracellular stiffening during chemotherapeutic drug exposure agree with earlier studies on the exposure of K562 and Jurkat leukaemia cells to daunorubicin [13], treatment of PC-3 pancreatic cancer cells with disulfiram, paclitaxel, tomatine, and valproic acid [14] and the effects of Cisplatin and docetaxel on prostate cancer cells (PNTIA and 22Rv1) [15].

The stiffening of cancer cells during exposure to paclitaxel is attributed to alterations in the cytoskeletal structure since the treatment targets the cytoskeleton to induce cancer cell death. Paclitaxel promotes the formation and stabilisation of microtubules, reducing the depolymerisation and dynamic reorganisation of the cytoskeleton required for cell division and proliferation [16–18]. As a result, the cytoskeletal density increases. The stabilised microtubules push the actin cortex outward, leading to cortical strain-hardening [32]. The increase in cytoskeletal density and the strain hardening of the cortex could explain the increase in intracellular stiffness of the paclitaxel-treated cells in the current study. Future studies will confirm whether the observed increase in the intracellular stiffness on exposure to paclitaxel is mainly due to alterations in the cytoskeleton. These studies will measure total cytoskeletal content through western blotting and immunofluorescence experiments and inhibition of the cytoskeletal components using drugs such as nocodazole, cytochalasin D, and blebbistatin.

### 4.2 Cell morphology

The morphometric results show that the paclitaxel-treated cells were significantly larger in cell area and perimeter than the untreated controls after 24 and 48 hours in 2D environments. The increased exposure time to paclitaxel of 48 versus 24 hours led to a significant increase in the perimeter (but not the cell area) of paclitaxel-treated cells, whereas the untreated cells did not change the size between 24 and 48 hours. A size increase due to paclitaxel exposure was not observed for cells in 3D environments.

The morphological results for cells in 2D agree with several previous studies demonstrating that cancer cells increase in size due to chemotherapeutic treatment [33–35]. When exposed to paclitaxel, the enlarging of cells can be attributed to the increase in cytoskeletal density based on paclitaxel-promoted formation and stabilisation of microtubules, leading to a reduction in their depolymerisation and dynamics [16–18]. Another mechanism of increasing cell size may be forming multiple nuclei within a single cell. Multi-nucleation was observed in the current study (see supplemental Figure S1) and ascribed to the inhibition of cell division by paclitaxel [34, 36].

The shape analysis results show that the cell shapes were well distributed among all the shape mode groups in the different treatment conditions except for group 1 in paclitaxel-treated and untreated cells at 48 hours and the untreated cells at 24 hours in 3D. No statistically significant differences were observed in the distribution of cell shapes across the different treatment conditions (considering Bonferroni adjusted results), indicating that paclitaxel treatment had no significant effect on the heterogeneity of cell shapes or that the stiffness changes observed may be insufficient to alter the shape of cells substantially both in 2D and 3D. More research with a larger sample size and long-term drug treatment are required to confirm these results. While the shape mode analysis in the current study was an explorative investigation into possible additional cellular biomarkers for chemotherapeutic drug response, knowledge of cell shape distribution will be useful for future research in this area.

The differences observed in the intracellular stiffness, morphology and distribution of cellular shapes between the cells in 2D and 3D environments can be attributed to the differences in geometry. Cells and the cytoskeleton are more planar in 2D than in 3D where they are more isotropic due to fewer geometrical constraints when attaching to the substrates [37].

## 5. Conclusion

This study has shown that paclitaxel treatment affects the intracellular stiffness of OSCC cells in both 2D and 3D environments and their morphology (cell area and perimeter when in 2D and but not heterogeneity of shape modes). This study used a clonal cell line derived from South African OSCC patients; however, future studies with primary human OSCC cells are required to ascertain the observed changes in the cellular mechanics. Additionally, the OSCC cells in this study were treated with paclitaxel for up to 48 hours. Prolonged exposure to chemotherapeutic drugs in vitro will help characterise the long-term effects of chemotherapeutic drugs on the physical properties of OSCC cell lines and primary cells. These physical characteristics of OSCC cells may complement genetic and molecular markers to assess the effectiveness of chemotherapy and onset of chemoresistance to improve the treatment success in OSCC patients.

## Acknowledgements

We thank the following colleagues from the University of Cape Town: Prof Iqbal Parker from the Department of Integrative Biomedical Sciences for donating WHOC1 cells for this study, Mrs Helen Ilsley from the Cardiovascular Research Unit for technical assistance with cell culturing and histochemistry, and Ms Susan Cooper of the Confocal and Light Microscope Imaging Facility for technical assistance with microscopic imaging. We also thank Prof Denis Wirtz from Johns Hopkins University, Baltimore, Maryland, USA, for providing the visually aided morpho-phenotyping image recognition (VAMPIRE) software.

## Data availability

Data supporting the results presented in this article are available on the University of Cape Town’s institutional data repository (ZivaHub) under https://doi.org/10.25375/uct.17693510 as Kiwanuka M, Higgins G, Ngcobo S, Nagawa J, Lang DM, Zaman MH, Davies NH, Franz T. Data for “Effect of paclitaxel treatment on cellular mechanics and morphology of oesophageal squamous cell carcinoma of South African origin in 2D and 3D environments”. Cape Town, ZivaHub, 2022, DOI: 10.25375/uct.17693510.

