## Supplemental figure S1 for "Effect of paclitaxel treatment on cellular mechanics and morphology of human oesophageal squamous cell carcinoma in 2D and 3D environments"

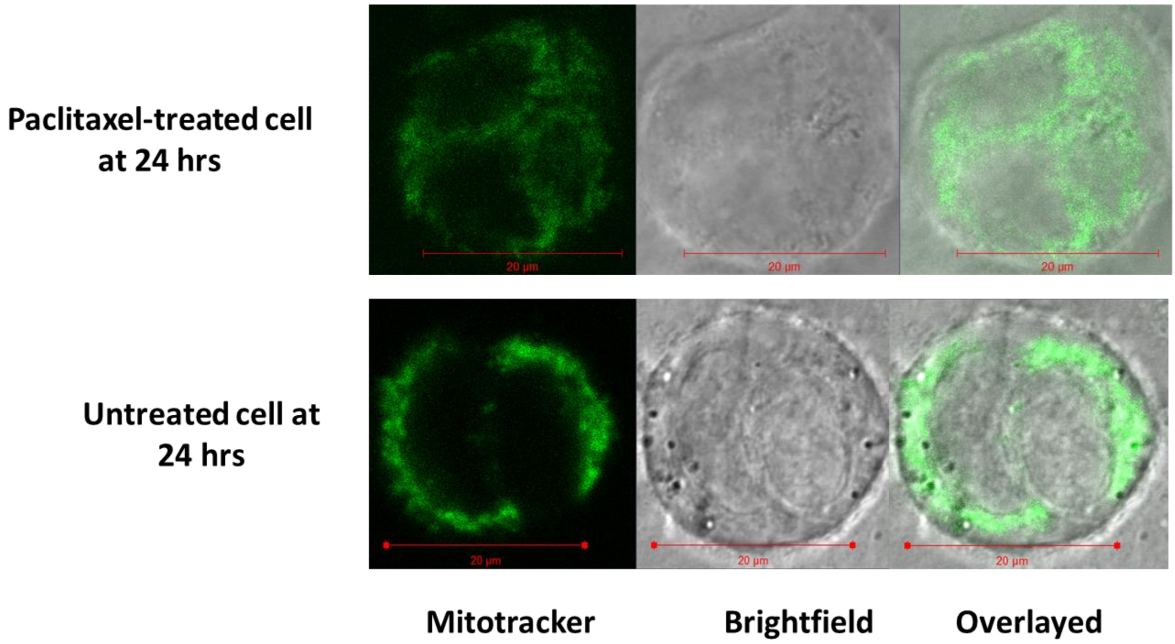


Figure S1: Multinucleated cells. Representative fluorescence, brightfield, and overlay images of paclitaxel-treated and untreated single WHCO1 cells showing the multiple nuclei in cells observed after 24 hours in 3D environments.
