## Supplemental data for "Effect of paclitaxel treatment on cellular mechanics and morphology of human oesophageal squamous cell carcinoma in 2D and 3D environments"

### 1 Microrheology data on cells in 2D environments

Table S 1: Summaries of mean MSDs and standard deviations of mitochondrial fluctuations from 79 individual WHCO1 cells in 2D environments at 24 and 48 hours of treatment. PTX and UNT stand for Paclitaxel-treated and untreated cells, respectively. The MSDs are $\mu m^2$. MSDs at $\tau = 0.7$ s were transformed using a logarithmic function, whereas $\tau = 6.3$, 7.7, and 9.1 s were transformed using a square root function, respectively, after failing the normality test.

| **Delay (s)** | **Mean (M)** | | | | | | **Standard Deviation (SD)** | | | | | |
|---|---|---|---|---|---|---|---|---|---|---|---|---|
| | **PTX** | | **UNT** | | **ETH** | | **PTX** | | **UNT** | | **ETH** | |
| | ***24*** | ***48*** | ***24*** | ***48*** | ***24*** | ***48*** | ***24*** | ***48*** | ***24*** | ***48*** | ***24*** | ***48*** |
| **0.14** | .003 | .003 | .003 | .003 | .003 | .003 | .0002 | .0004 | .0003 | .0002 | .0003 | .0004 |
| **0.7** | -2.1 | -2.1 | -2.1 | -2.1 | -2.2 | -2.3 | .1009 | .0494 | .0673 | .1249 | .1261 | .0886 |
| **3.5** | 0.04 | 0.03 | 0.03 | 0.03 | 0.03 | 0.01 | .0142 | .0054 | .0095 | .0129 | .0138 | .0067 |
| **6.3** | .241 | .224 | .243 | .235 | .209 | .156 | .0540 | .0260 | .0404 | .0492 | .0668 | .0434 |
| **7.7** | .263 | .241 | .268 | .257 | .231 | .172 | .061 | .031 | .049 | .054 | .078 | .051 |
| **9.1** | .284 | .261 | .290 | .277 | .252 | .188 | .067 | .034 | .059 | .059 | .091 | .060 |

Table S 2: Summaries of mean fluidity and standard deviations of mitochondrial fluctuations from 79 individual WHCO1 cells in 2D environments at 24 and 48 hours of treatment. PTX and UNT stand for Paclitaxel-treated and untreated cells, respectively. Fluidity values at $\tau = 6.3$ and 9.1 s were transformed using a logarithmic and square root function, respectively, after failing the normality test.

| **Delay (s)** | **Mean (M)** | | | | | | **Standard Deviation (SD)** | | | | | |
|---|---|---|---|---|---|---|---|---|---|---|---|---|
| | **UNT** | | **ETH** | | **PTX** | | **UNT** | | **ETH** | | **PTX** | |
| | ***24*** | ***48*** | ***24*** | ***48*** | ***24*** | ***48*** | ***24*** | ***48*** | ***24*** | ***48*** | ***24*** | ***48*** |
| **0.28** | 0.43 | 0.40 | 0.41 | 0.42 | 0.31 | 0.20 | 0.15 | 0.07 | 0.11 | 0.14 | 0.12 | 0.09 |
| **0.7** | 0.73 | 0.72 | 0.73 | 0.71 | 0.61 | 0.38 | 0.18 | 0.10 | 0.13 | 0.20 | 0.19 | 0.20 |
| **2.1** | 0.92 | 0.91 | 0.96 | 0.85 | 0.85 | 0.68 | 0.10 | 0.07 | 0.12 | 0.13 | 0.15 | 0.21 |
| **3.5** | 0.91 | 0.88 | 0.96 | 0.88 | 0.91 | 0.82 | 0.11 | 0.11 | 0.11 | 0.12 | 0.15 | 0.18 |
| **4.9** | 0.91 | 0.83 | 0.93 | 0.84 | 0.95 | 0.90 | 0.11 | 0.14 | 0.19 | 0.15 | 0.23 | 0.20 |
| **6.3** | -0.05 | -0.08 | -0.04 | -0.06 | -0.04 | -0.03 | 0.07 | 0.08 | 0.12 | 0.07 | 0.10 | 0.09 |
| **9.1** | 0.91 | 0.88 | 0.92 | 0.95 | 0.93 | 0.95 | 0.10 | 0.10 | 0.22 | 0.10 | 0.18 | 0.21 |

Table S 3: Two-way ANOVA results summarise the significance of the interaction between treatment intervention (untreated and Paclitaxel-treated cells) and duration of treatment (24 and 48 hours) in MSDs and fluidity at the different delay times for the cells in 2D.

| **Delay (s)** | **MSDs** | | | **Delay (s)** | **Fluidity (α)** | | |
|---|---|---|---|---|---|---|---|
| | ***F (2, 73)*** | ***p*** | $\eta_p^2$ | | ***F (2, 73)*** | ***p*** | $\eta_p^2$ |
| **0.14** | 0.554 | 0.577 | 0.015 | **0.28** | 1.831 | 0.168 | 0.048 |
| **0.7** | 2.901 | 0.061 | 0.074 | **0.7** | 3.758 | 0.028 | 0.093 |
| **2.1** | 1.943 | 0.151 | 0.051 | **2.1** | 2.325 | 0.105 | 0.060 |
| **3.5** | 1.511 | 0.228 | 0.040 | **3.5** | 0.292 | 0.748 | 0.008 |
| **4.9** | 2.596 | 0.081 | 0.066 | **4.9** | 0.150 | 0.861 | 0.004 |
| **6.3** | 1.511 | 0.228 | 0.040 | **6.3** | 0.382 | 0.684 | 0.010 |
| **7.7** | 1.343 | 0.267 | 0.035 | **7.7** | 0.255 | 0.775 | 0.007 |
| **9.1** | 1.173 | 0.315 | 0.031 | **9.1** | 0.309 | 0.735 | 0.008 |

Table S 4: Pairwise comparisons of MSDs of Paclitaxel-treated (PTX or P), ethanol-treated (ETH or E), and untreated (UNT or U) in 2D environments at 24 and 48 hours of treatment. For example, PTX (24 Vs. 48) compares MSDs for Paclitaxel-treated cells at 24 and 48 hours.

| **Delay (s)** | **P-values** | | | | | | | | |
|---|---|---|---|---|---|---|---|---|---|
| | **UNT** ***(24 Vs 48)*** | **ETH** ***(24 Vs 48)*** | **PTX** ***(24 Vs 48)*** | **24** | | | **48** | | |
| | | | | ***(U Vs E)*** | ***(E Vs P)*** | ***(P Vs U)*** | ***(U Vs E)*** | ***(E Vs P)*** | ***(P Vs U)*** |
| **0.14** | 0.625 | 0.362 | 0.631 | 1.000 | 1.000 | 1.000 | 0.109 | 0.411 | 1.000 |
| **0.7** | 0.382 | 0.548 | 0.006 | 1.000 | 0.078 | 0.031 | 0.580 | <.0005 | <.0005 |
| **3.5** | 0.245 | 0.984 | 0.014 | 1.000 | 0.277 | 0.125 | 1.000 | <.0005 | 0.002 |
| **6.3** | 0.335 | 0.683 | 0.007 | 1.000 | 0.271 | 0.249 | 1.000 | <.0005 | 0.001 |
| **7.7** | 0.324 | 0.650 | 0.008 | 1.000 | 0.348 | 0.370 | 1.000 | 0.001 | 0.003 |
| **9.1** | 0.330 | 0.646 | 0.012 | 1.000 | 0.477 | 0.571 | 1.000 | 0.002 | 0.009 |

Table S 5: Pairwise comparisons of fluidity for Paclitaxel-treated (PTX or P), ethanol-treated (ETH or E), and untreated (UNT, or U) in 2D environments at 24 and 48 hours of treatment.

| **Delay (s)** | **P-values** | | | | | | | | |
|---|---|---|---|---|---|---|---|---|---|
| | **UNT** ***(24 Vs 48)*** | **ETH** ***(24 Vs 48)*** | **PTX** ***(24 Vs 48)*** | **24** | | | **48** | | |
| | | | | ***(U Vs E)*** | ***(E Vs P)*** | ***(P Vs U)*** | ***(U Vs E)*** | ***(E Vs P)*** | ***(P Vs U)*** |
| **0.28** | 0.424 | 0.755 | 0.017 | 1.000 | 0.177 | 0.026 | 1.000 | <.0005 | <.0005 |
| **0.7** | 0.831 | 0.754 | 0.001 | 1.000 | 0.290 | 0.197 | 1.000 | <.0005 | <.0005 |
| **2.1** | 0.832 | 0.068 | 0.003 | 1.000 | 0.178 | 0.500 | 0.838 | 0.010 | <.0005 |
| **3.5** | 0.497 | 0.191 | 0.117 | 1.000 | 1.000 | 1.000 | 1.000 | 0.858 | 0.868 |
| **4.9** | 0.204 | 0.186 | 0.510 | 1.000 | 1.000 | 1.000 | 1.000 | 1.000 | 0.729 |
| **6.3** | 0.319 | 0.610 | 0.802 | 1.000 | 1.000 | 1.000 | 1.000 | 1.000 | 0.407 |
| **9.1** | 0.536 | 0.663 | 0.839 | 1.000 | 1.000 | 1.000 | 0.700 | 1.000 | 0.693 |

#### 2 Microrheology data of cells in 3D matrices

Table S 6: Summarises mean MSDs and standard deviations of mitochondrial fluctuations from 63 individual WHCO1 cells embedded in 3D environments at 24 and 48 hours of treatment. PTX and UNT stand for Paclitaxel-treated and untreated cells, respectively. The MSDs are $\mu m^2$.

| Delay (s) | Mean (M) | | | | Standard Deviation (SD) | | | |
|---|---|---|---|---|---|---|---|---|
| | PTX | | UNT | | PTX | | UNT | |
| | *24* | *48* | *24* | *48* | *24* | *48* | *24* | *48* |
| **0.15** | 0.0233 | 0.0268 | 0.0241 | 0.0233 | 0.0063 | 0.0033 | 0.0042 | 0.0034 |
| **0.3** | 0.0261 | 0.0313 | 0.0283 | 0.0264 | 0.0069 | 0.0043 | 0.0047 | 0.0037 |
| **0.6** | 0.0294 | 0.0372 | 0.0345 | 0.0309 | 0.0075 | 0.0063 | 0.0051 | 0.0041 |
| **0.9** | 0.0326 | 0.0432 | 0.0402 | 0.0350 | 0.0081 | 0.0086 | 0.0059 | 0.0048 |
| **1.2** | 0.0357 | 0.0485 | 0.0458 | 0.0392 | 0.0092 | 0.0112 | 0.0068 | 0.0058 |
| **1.5** | 0.0382 | 0.0536 | 0.0510 | 0.0430 | 0.0097 | 0.0120 | 0.0078 | 0.0069 |
| **1.8** | 0.0406 | 0.0586 | 0.0560 | 0.0466 | 0.0103 | 0.0165 | 0.0090 | 0.0077 |
| **2.1** | 0.0431 | 0.0633 | 0.0608 | 0.0504 | 0.0113 | 0.0192 | 0.0102 | 0.0087 |
| **2.4** | 0.0454 | 0.0681 | 0.0655 | 0.0542 | 0.0120 | 0.2193 | 0.0114 | 0.0098 |
| **2.7** | 0.0477 | 0.0729 | 0.0702 | 0.0580 | 0.0128 | 0.0247 | 0.0127 | 0.0110 |
| **3** | 0.0499 | 0.0777 | 0.0747 | 0.0617 | 0.0136 | 0.0275 | 0.0140 | 0.0121 |
| **3.6** | 0.0540 | 0.0867 | 0.0837 | 0.0689 | 0.0153 | 0.0330 | 0.0164 | 0.0145 |
| **4.2** | 0.0580 | 0.0952 | 0.0921 | 0.0765 | 0.0171 | 0.0382 | 0.0189 | 0.0174 |
| **4.8** | 0.0619 | 0.1038 | 0.1005 | 0.0841 | 0.0187 | 0.0435 | 0.0216 | 0.0207 |
| **6** | 0.0690 | 0.1195 | 0.1167 | 0.0994 | 0.0217 | 0.0537 | 0.0277 | 0.0275 |
| **9.9** | 0.0848 | 0.1543 | 0.1540 | 0.1381 | 0.0300 | 0.0768 | 0.0416 | 0.0464 |

Table S 7: Summarises mean fluidity and standard deviations of mitochondrial fluctuations from 63 individual WHCO1 cells embedded in 3D environments at 24 and 48 hours of treatment. PTX and UNT stand for Paclitaxel-treated and untreated cells, respectively. Fluidity at $\tau$ = 1.5 s was transformed using a square root function after failing the normality test.

| Delay (s) | Mean (M) | | | | Standard Deviation (SD) | | | |
|---|---|---|---|---|---|---|---|---|
| | PTX | | UNT | | PTX | | UNT | |
| | *24* | *48* | *24* | *48* | *24* | *48* | *24* | *48* |
| **0.3** | 0.1666 | 0.2175 | 0.2365 | 0.1803 | 0.0431 | 0.0773 | 0.0395 | 0.0378 |
| **0.6** | 0.1941 | 0.2661 | 0.3245 | 0.2483 | 0.0850 | 0.1091 | 0.0797 | 0.0798 |
| **0.9** | 0.2675 | 0.3671 | 0.4012 | 0.3264 | 0.0894 | 0.1289 | 0.1044 | 0.0997 |
| **1.2** | 0.3009 | 0.3848 | 0.4633 | 0.3934 | 0.1243 | 0.1378 | 0.1012 | 0.0943 |
| **1.5** | 0.5728 | 0.6406 | 0.6856 | 0.6454 | 0.1240 | 0.1343 | 0.0653 | 0.1014 |
| **1.8** | 0.3397 | 0.4959 | 0.5150 | 0.4572 | 0.1534 | 0.1397 | 0.1096 | 0.1481 |
| **2.1** | 0.3871 | 0.4815 | 0.5289 | 0.5153 | 0.1629 | 0.1584 | 0.1132 | 0.1357 |
| **2.4** | 0.3804 | 0.5125 | 0.5530 | 0.5388 | 0.1727 | 0.1288 | 0.1055 | 0.1620 |
| **2.7** | 0.4144 | 0.5434 | 0.5852 | 0.5831 | 0.1325 | 0.1828 | 0.0918 | 0.1567 |
| **3** | 0.4509 | 0.5608 | 0.5664 | 0.5748 | 0.1497 | 0.1408 | 0.1215 | 0.1321 |
| **3.6** | 0.4501 | 0.5772 | 0.6113 | 0.6231 | 0.1973 | 0.1400 | 0.0863 | 0.1691 |
| **4.2** | 0.5033 | 0.5387 | 0.6143 | 0.6160 | 0.1547 | 0.1557 | 0.1229 | 0.2098 |
| **4.8** | 0.4386 | 0.6386 | 0.6388 | 0.7123 | 0.1611 | 0.1832 | 0.1258 | 0.2041 |
| **6** | 0.5235 | 0.5965 | 0.6882 | 0.7222 | 0.2534 | 0.2443 | 0.1458 | 0.2334 |
| **9.9** | 0.4758 | 0.6995 | 0.7084 | 0.7540 | 0.4010 | 0.4908 | 0.1991 | 0.2462 |

Table S 8: Two-way ANOVA results summarise the significance of the interaction between treatment intervention (untreated and Paclitaxel-treated cells) and duration of treatment (24 and 48 hours) in MSDs and fluidity at the different delay times for the cells in 3D.

| **Delay (s)** | **MSDs** | | | **Fluidity (α)** | | |
|---|---|---|---|---|---|---|
| | ***F (1, 59)*** | ***p*** | $\eta_p^2$ | ***F (1, 59)*** | ***p*** | $\eta_p^2$ |
| **0.15** | 2.931 | 0.092 | 0.047 | | | |
| **0.3** | 6.270 | 0.015 | 0.096 | 17.908 | <.0005 | 0.233 |
| **0.6** | 12.694 | 0.001 | 0.177 | 10.551 | 0.002 | 0.152 |
| **0.9** | 18.390 | < .0005 | 0.238 | 10.364 | 0.002 | 0.149 |
| **1.2** | 19.893 | < .0005 | 0.252 | 6.522 | 0.013 | 0.100 |
| **1.5** | 21.878 | < .0005 | 0.271 | 3.697 | 0.059 | 0.060 |
| **1.8** | 23.484 | < .0005 | 0.285 | 8.666 | 0.005 | 0.128 |
| **2.1** | 23.011 | < .0005 | 0.281 | 2.052 | 0.157 | 0.034 |
| **2.4** | 23.011 | < .0005 | 0.281 | 3.616 | 0.062 | 0.058 |
| **2.7** | 22.734 | < .0005 | 0.278 | 3.332 | 0.073 | 0.053 |
| **3** | 22.393 | < .0005 | 0.275 | 1.992 | 0.163 | 0.033 |
| **3.6** | 21.896 | < .0005 | 0.271 | 1.977 | 0.165 | 0.032 |
| **4.2** | 20.390 | < .0005 | 0.257 | 0.162 | 0.688 | 0.003 |
| **4.8** | 19.268 | < .0005 | 0.246 | 2.117 | 0.151 | 0.035 |
| **6** | 16.773 | < .0005 | 0.221 | 0.114 | 0.737 | 0.002 |
| **9.9** | 12.123 | 0.001 | 0.170 | 0.991 | 0.324 | 0.017 |

Table S 9: Pairwise comparisons of MSDs of Paclitaxel-treated (PTX) and untreated (UNT) in 3D environments at 24 and 48 hours of treatment. For example, PTX (24 Vs. 48) compares MSDs for Paclitaxel-treated cells at 24 and 48 hours.

| **Delay (s)** | **P-values** | | | |
|---|---|---|---|---|
| | **PTX** ***(24 Vs 48)*** | **UNT** ***(24 Vs 48)*** | **24** ***(UNT Vs PTX)*** | **48** ***(UNT Vs PTX)*** |
| **0.15** | 0.056 | 0.676 | 0.596 | 0.087 |
| **0.3** | 0.015 | 0.333 | 0.195 | 0.036 |
| **0.6** | 0.001 | 0.107 | 0.011 | 0.016 |
| **0.9** | < .0005 | 0.042 | 0.001 | 0.007 |
| **1.2** | < .0005 | 0.030 | < .0005 | 0.010 |
| **1.5** | < .0005 | 0.023 | < .0005 | 0.010 |
| **1.8** | < .0005 | 0.018 | < .0005 | 0.010 |
| **2.1** | < .0005 | 0.020 | < .0005 | 0.014 |
| **2.4** | < .0005 | 0.023 | < .0005 | 0.017 |
| **2.7** | < .0005 | 0.027 | < .0005 | 0.021 |
| **3** | < .0005 | 0.031 | < .0005 | 0.024 |
| **3.6** | < .0005 | 0.038 | < .0005 | 0.033 |
| **4.2** | < .0005 | 0.056 | < .0005 | 0.051 |
| **4.8** | < .0005 | 0.076 | < .0005 | 0.068 |
| **6** | < .0005 | 0.132 | < .0005 | 0.133 |
| **9.9** | < .0005 | 0.346 | < .0005 | 0.411 |

Table S 10: Pairwise comparisons of fluidity for Paclitaxel-treated (PTX) and untreated (UNT) in 3D environments at 24 and 48 hours of treatment.

| **Delay (s)** | **P-values** | | | |
|---|---|---|---|---|
| | **PTX** *(24 Vs 48)* | **UNT** *(24 Vs 48)* | **24** *(UNT Vs PTX)* | **48** *(UNT Vs PTX)* |
| **0.3** | 0.008 | 0.002 | < .0005 | 0.071 |
| **0.6** | 0.035 | 0.018 | < .0005 | 0.626 |
| **0.9** | 0.014 | 0.048 | < .0005 | 0.351 |
| **1.2** | 0.061 | 0.095 | < .0005 | 0.858 |
| **1.5** | 0.105 | 0.297 | 0.002 | 0.914 |
| **1.8** | 0.005 | 0.250 | < .0005 | 0.508 |
| **2.1** | 0.091 | 0.794 | 0.003 | 0.575 |
| **2.4** | 0.022 | 0.788 | < .0005 | 0.671 |
| **2.7** | 0.017 | 0.966 | < .0005 | 0.492 |
| **3** | 0.041 | 0.866 | 0.010 | 0.809 |
| **3.6** | 0.038 | 0.834 | 0.002 | 0.487 |
| **4.2** | 0.565 | 0.977 | 0.032 | 0.254 |
| **4.8** | 0.003 | 0.220 | < .0005 | 0.292 |
| **6** | 0.391 | 0.670 | 0.021 | 0.180 |
| **9.9** | 0.092 | 0.711 | 0.035 | 0.705 |

#### 3 Morphological analysis

Table S 11: Summaries of mean and standard deviations for the area, perimeter, circularity, and aspect ratio from WHCO1 cells in 2D and 3D environments at 24 and 48 hours of treatment. PTX and UNT stand for Paclitaxel-treated and untreated cells, respectively.

| Dependent variable | Time (hours) | Intervention | Dimension | Mean (M) | Standard deviation (SD) |
|---|---|---|---|---|---|
| **Area** | 24 | PTX | 2D | 46.553 | 8.921 |
| | | | 3D | 77.061 | 14.768 |
| | | UNT | 2D | 30.973 | 5.936 |
| | | | 3D | 82.477 | 15.261 |
| | 48 | PTX | 2D | 55.709 | 10.676 |
| | | | 3D | 84.270 | 16.843 |
| | | UNT | 2D | 35.078 | 6.722 |
| | | | 3D | 91.186 | 15.615 |
| **Perimeter** | 24 | PTX | 2D | 1.100 | 0.143 |
| | | | 3D | 1.434 | 0.175 |
| | | UNT | 2D | 1.010 | 0.123 |
| | | | 3D | 1.670 | 0.258 |
| | 48 | PTX | 2D | 1.157 | 0.141 |
| | | | 3D | 1.512 | 0.171 |
| | | UNT | 2D | 0.964 | 0.118 |
| | | | 3D | 1.658 | 0.252 |
| **Circularity** | 24 | PTX | 2D | 0.510 | 0.104 |
| | | | 3D | 0.477 | 0.097 |
| | | UNT | 2D | 0.406 | 0.083 |
| | | | 3D | 0.380 | 0.073 |
| | 48 | PTX | 2D | 0.415 | 0.084 |
| | | | 3D | 0.464 | 0.071 |
| | | UNT | 2D | 0.476 | 0.097 |
| | | | 3D | 0.430 | 0.090 |
| **Aspect Ratio** | 24 | PTX | 2D | 1.349 | 0.113 |
| | | | 3D | 1.142 | 0.096 |
| | | UNT | 2D | 1.451 | 0.122 |
| | | | 3D | 1.165 | 0.116 |
| | 48 | PTX | 2D | 1.580 | 0.133 |
| | | | 3D | 1.109 | 0.060 |
| | | UNT | 2D | 1.448 | 0.122 |
| | | | 3D | 1.165 | 0.107 |

Table S 12: Pairwise comparisons for mean area, perimeter, circularity, and aspect ratio for Paclitaxel-treated (PTX) and untreated (UNT) in 2D and 3D environments at 24 and 48 hours

| Dependent variable | Interventions | | Mean Differences | P-Value |
|---|---|---|---|---|
| **Area** | **2D** | **24** | **PTX** *(2D V 3D)* | < .0005 |
| | | | **UNT** *(2D v 3D)* | < .0005 |
| | | | *PTX v UNT* | 0.001 |
| | | | *PTX v ETH* | < .0005 |
| | | | *ETH v UNT* | 1.000 |
| | | **48** | **PTX** *(3D v 2D)* | < .0005 |
| | | | **UNT** *(2D v 3D)* | < .0005 |
| | | | *PTX v UNT* | < .0005 |
| | | | *PTX v ETH* | < .0005 |
| | | | *ETH v UNT* | 1.000 |
| | **3D** | **24** | *PTX v UNT* | 0.399 |
| | | **48** | *PTX v UNT* | 0.330 |
| | **PTX** | **2D** | *48 v 24* | 0.075 |
| | | **3D** | *48 v 24* | 0.310 |
| | **ETH** | **2D** | *48 v 24* | 0.721 |
| | **UNT** | **2D** | *48 v 24* | 0.308 |
| | | **3D** | *48 v 24* | 0.176 |
| **Perimeter** | **2D** | **24** | **PTX** *(2D V 3D)* | 0.001 |
| | | | **UNT** *(2D v 3D)* | < .0005 |
| | | | *PTX v UNT* | 0.737 |
| | | | *PTX v ETH* | 0.001 |
| | | | *ETH v UNT* | 0.012 |
| | | **48** | **PTX** *(3D v 2D)* | 0.190 |
| | | | **UNT** *(2D v 3D)* | < .0005 |
| | | | *PTX v UNT* | < .0005 |
| | | | *PTX v ETH* | < .0005 |
| | | | *ETH v UNT* | 1.000 |
| | **3D** | **24** | *PTX v UNT* | 0.037 |
| | | **48** | *PTX v UNT* | 0.241 |
| | **PTX** | **2D** | *48 v 24* | 0.004 |
| | | **3D** | *48 v 24* | 0.530 |
| | **ETH** | **2D** | *48 v 24* | 0.142 |
| | **UNT** | **2D** | *48 v 24* | 0.512 |
| | | **3D** | *48 v 24* | 0.916 |
| **Circularity** | **2D** | **24** | **PTX** *(2D V 3D)* | 0.359 |
| | | | **UNT** *(2D v 3D)* | 0.436 |
| | | | *PTX v UNT* | < .0005 |
| | | | *PTX v ETH* | 0.142 |
| | | | *ETH v UNT* | < .0005 |
| | | **48** | **PTX** *(3D v 2D)* | 0.227 |
| | | | **UNT** *(2D v 3D)* | 0.175 |
| | | | *PTX v UNT* | 0.124 |
| | | | *PTX v ETH* | 0.952 |
| | | | *ETH v UNT* | 1.000 |
| | **3D** | **24** | *PTX v UNT* | 0.015 |
| | | **48** | *PTX v UNT* | 0.433 |
| | **PTX** | **2D** | *48 v 24* | 0.003 |
| | | **3D** | *48 v 24* | 0.765 |
| | **ETH** | **2D** | *48 v 24* | < .0005 |
| | **UNT** | **2D** | *48 v 24* | 0.005 |
| | | **3D** | *48 v 24* | 0.214 |

Table S12 continued

| Dependent variable | Interventions | | Mean Differences | P-Value |
|---|---|---|---|---|
| **Aspect Ratio** | **2D** | **24** | **PTX** *(2D V 3D)* | 0.095 |
| | | | **UNT** *(2D v 3D)* | 0.012 |
| | | | *PTX v UNT* | 0.855 |
| | | | *PTX v ETH* | 1.000 |
| | | | *ETH v UNT* | 0.208 |
| | | **48** | **PTX** *(3D v 2D)* | 0.001 |
| | | | **UNT** *(2D v 3D)* | 0.017 |
| | | | *PTX v UNT* | 0.609 |
| | | | *PTX v ETH* | 0.087 |
| | | | *ETH v UNT* | 0.728 |
| | **3D** | **24** | *PTX v UNT* | 0.870 |
| | | **48** | *PTX v UNT* | 0.714 |
| | **PTX** | **2D** | *48 v 24* | 0.038 |
| | | **3D** | *48 v 24* | 0.830 |
| | **ETH** | **2D** | *48 v 24* | 0.765 |
| | **UNT** | **2D** | *48 v 24* | 0.974 |
| | | **3D** | *48 v 24* | 0.997 |

Table S 13: VAMPIRE analysis results showing the abundance of cells in each condition corresponding to each shape mode. P_24, P_48, E_24, E_48, U_24, U_48 represent the number of Paclitaxel-treated cells at 24 and 48 hours, ethanol-treated cells at 24 and 48 hours, and untreated cells at 24 and 48 hours, respectively.

| **Shape Modes** | **2D** | | | | | | **3D** | | | |
|---|---|---|---|---|---|---|---|---|---|---|
| | **P_24** | **P_48** | **E_24** | **E_48** | **U_24** | **U_48** | **P_24** | **P_48** | **U_24** | **U_48** |
| **1** | 2 | 3 | 5 | 1 | 4 | 9 | 1 | 0 | 0 | 0 |
| **2** | 5 | 2 | 2 | 1 | 3 | 3 | 5 | 4 | 3 | 4 |
| **3** | 2 | 1 | 5 | 3 | 4 | 0 | 3 | 0 | 2 | 3 |
| **4** | 4 | 2 | 8 | 4 | 3 | 4 | 1 | 3 | 1 | 3 |
| **5** | 4 | 0 | 6 | 4 | 4 | 3 | 1 | 0 | 2 | 1 |
| **6** | 2 | 4 | 6 | 0 | 5 | 4 | 2 | 0 | 1 | 0 |
| **7** | 2 | 4 | 2 | 3 | 8 | 0 | 1 | 1 | 1 | 0 |
| **8** | 2 | 2 | 8 | 2 | 5 | 3 | 0 | 0 | 1 | 1 |
| **9** | 3 | 0 | 0 | 1 | 2 | 4 | 1 | 3 | 4 | 1 |
| **10** | 1 | 4 | 1 | 4 | 4 | 7 | 0 | 0 | 1 | 2 |
